# Calling pangenes from plant genome alignments confirms presence-absence variation

**DOI:** 10.1101/2023.01.03.520531

**Authors:** Bruno Contreras-Moreira, Shradha Saraf, Guy Naamati, Ana M. Casas, Sandeep S. Amberkar, Paul Flicek, Andrew R. Jones, Sarah Dyer

**Affiliations:** European Molecular Biology Laboratory, European Bioinformatics Institute, Hinxton, UK; Estación Experimental Aula Dei-CSIC, Zaragoza, 50059, Spain; Institute of Systems, Molecular and Integrative Biology, University of Liverpool, UK

## Abstract

Consistent gene annotation in crops is becoming harder as genomes for new cultivars are frequently published. Gene sets from recently sequenced accessions have different gene identifiers to those on the reference accession, and might be of higher quality due to technical advances. For these reasons there is a need to define pangenes, which represent all known syntenic orthologues for a gene model and can be linked back to the original annotation sources. A pangene set effectively summarizes our current understanding of the coding potential of a crop and can be used to inform gene model annotation in new cultivars. Here we present an approach (get_pangenes) to identify and analyze pangenes that is not biased towards the reference annotation. The method involves computing Whole Genome Alignments (WGA), which are used to estimate gene model overlaps. After a benchmark on *Arabidopsis*, rice, wheat and barley datasets, we find that minimap2 performs better than the GSAlign WGA algorithm. Our results show that pangenes recapitulate known phylogeny-based orthologies while adding extra core gene models in rice. More importantly, get_pangenes can also produce clusters of genome segments (gDNA) that overlap with gene models annotated in other cultivars. By lifting-over CDS sequences, gDNA clusters can help refine gene models across individuals and confirm or reject observed gene Presence-Absence Variation. A collection of flowering-related genes from the barley pangenome are discussed in detail. Documentation and source code are available at https://github.com/Ensembl/plant-scripts.

## INTRODUCTION

For a growing number of crops and plants there are now multiple genome assemblies available in public repositories. These data are driving the analysis of the pangenome, the union of all known genomes of a species. For instance, recently published pangenome reports include staple crops wheat and barley (Jayakodi et al., 2020; Walkowiak et al., 2020). While these efforts have greatly advanced our understanding of the variability of genomes within species, they have also prompted a new class of problems, those related to annotating and naming genes across cultivars. Different strategies are possible. For instance, the barley pangenome consortium lifted-over gene models from three genotypes (Morex, Barke, HOR10350) to all other assemblies. This procedure biases the gene space to that of the reference cultivars. In contrast, in other species, fresh gene annotations have been produced for different individuals or sampled populations (Gordon et al., 2017). In this case, care should be taken to follow the same annotation protocols throughout to avoid inflating the number of population-specific genes (Weisman et al., 2022) or to conserve gene identifiers. In this context it is useful to define a pangene, a gene model or allele found in some or all individuals of a species in a similar genomic location. A pangene should integrate additional naming schemes, e.g. so that a cluster of gene models can share a common identifier that links back to their original gene identifiers. A pangene set defines our current understanding of the total coding potential of a species and can assist in gene model curation, by providing a pool of possible gene models for assessment.

Pangenes can be produced by a variety of approaches, such as iterative mapping and assembly (Golicz et al., 2015), local alignments of nucleotide sequences (Contreras-Moreira et al., 2017), molecular phylogenies of chromosome-sorted proteins (Lovell et al., 2022) or as a secondary product of genome graphs (Sheikhizadeh et al., 2016; Li et al., 2020; Guarracino et al., 2022). Whatever the approach, a common use case for pangenes is to capture Presence Absence Variation (PAV) at the gene level. However, previous work has observed that absent gene models are rarely caused by complete sequence deletions; instead, they might not be expressed in certain conditions or the underlying genomic regions might contain genetic variants such that the criteria for calling a gene model are not satisfied. For instance, sequence variants in introns or splice sites can reduce evidence for a gene model (Lovell et al., 2021). Mapping transcript isoforms from orthologous loci is also a useful way to determine whether a gene model is intact (Kirilenko et al., 2023).

Here we present an approach to identify and analyze pangenes in sets of plant genomes which can explicitly confirm or reject PAVs by lifting-over viable gene models on candidate genomic segments. This approach requires computing pairwise Whole Genome Alignments (WGAs), which are then used to estimate gene model overlaps across individuals. Finally, pairs of overlapping genes are iteratively merged to produce pangene clusters. The algorithm produces pangene clusters that are not biased towards the reference annotation and that can optionally be used to refine individual gene model annotation with information from all cultivars. We benchmark this approach on diverse datasets that cover monocots and dicots, as well as small and large genomes.

## MATERIALS AND METHODS

### Genome sequences and genesets

A total of four datasets were used in this study (Arabidopsis ACK2, rice3, chr1wheat10 and barley20) of increasing size and complexity. They are listed in **Table 1**, where ACK stands for the Ancestral Crucifer Karyotype (Lysak et al., 2016) and chr1wheat10 for chromosome 1 in ten different hexaploid wheats. Note that ACK2 and rice3 include different species (*Arabidopsis lyrata* and *Oryza nivara*); the others include cultivars of the same species. The ranges of Average Nucleotide Identity (ANI) among genomes in each dataset are indicated in **Table 3**.

**Table 1.** Datasets used in this study. Geneset sources correspond to Ensembl Plants releases (EP) and the barley gene annotation from IPK (Mascher, M, 2020). Wheat genotypes are all *Triticum aestivum*. Barley genotypes are *Hordeum vulgare* landraces and cultivars, except B1K-04-12, which is *H. vulgare* subsp. *spontaneum*.

| dataset | species / genotype | INSDC accession | geneset | size (MB) | #genes |
| --- | --- | --- | --- | --- | --- |
| ACK2 | *Arabidopsis thaliana* | GCA_000001735.1 | EP52 | 116 | 27,655 |
|  | *Arabidopsis lyrata* | GCA_000004255.1 | EP52 | 200 | 32,667 |
| rice3 | *Oryza sativa* Nipponbare | GCA_001433935.1 | EP54 | 363 | 37,960 |
|  | *Oryza sativa* indica 93-11 | GCA_000004655.2 | EP54 | 414 | 40,745 |
|  | *Oryza nivara* IRGC:100I897 | GCA_000576065.1 | EP54 | 327 | 36,313 |
| chr1wheat10 | Chinese Spring | GCA_900519105.1 | EP52 | 1,736 | 18,017 |
|  | Arinalrfor | GCA_903993985.1 | EP52 | 1,725 | 13,542 |
|  | Jagger | GCA_903993795.1 | EP52 | 1,740 | 17,810 |
|  | Julius | GCA_903994195.1 | EP52 | 1,729 | 17,669 |
|  | Lancer | GCA_903993975.1 | EP52 | 1,721 | 17,951 |
|  | Landmark | GCA_903995565.1 | EP52 | 1,738 | 17,792 |
|  | Mace | GCA_903994175.1 | EP52 | 1,713 | 17,724 |
|  | Sy Mattis | GCA_903994185.1 | EP52 | 1,718 | 17,941 |
|  | Norin61 | GCA_904066035.1 | EP52 | 1,726 | 18,091 |
|  | Stanley | GCA_903994155.1 | EP52 | 1,737 | 17,767 |
| barley20 | MorexV2 | LR722616-LR7226<br>23 | IPK2020 | 4,210 | 46,294 |
|  | Akashinriki | ERS4201448 | IPK2020 | 4,401 | 44,446 |
|  | B1K-04-12 | ERS4201449 | IPK2020 | 4,142 | 44,566 |
|  | Barke | ERS4201450 | IPK2020 | 4,073 | 45,999 |
|  | Golden Promise | GCA_902500625.1 | IPK2020 | 3,946 | 42,464 |
|  | HOR10350 | ERS4201451 | IPK2020 | 4,086 | 45,810 |
|  | HOR13821 | ERS4201452 | IPK2020 | 4,324 | 44,714 |
|  | HOR13942 | ERS4201453 | IPK2020 | 4,249 | 44,718 |
|  | HOR21599 | ERS4201454 | IPK2020 | 4,344 | 44,456 |
|  | HOR3081 | ERS4201455 | IPK2020 | 4,201 | 45,146 |
|  | HOR3365 | ERS4201456 | IPK2020 | 4,722 | 47,588 |
|  | HOR7552 | ERS4201457 | IPK2020 | 4,228 | 44,641 |
|  | HOR8148 | ERS4201458 | IPK2020 | 4,212 | 45,026 |
|  | HOR9043 | ERS4201459 | IPK2020 | 4,270 | 45,028 |
|  | Hockett | ERS4201460 | IPK2020 | 4,201 | 46,450 |
|  | Igri | ERS4201461 | IPK2020 | 4,202 | 45,213 |
|  | OUN333 | ERS4201462 | IPK2020 | 4,392 | 44,699 |
|  | RGT Planet | ERS4201463 | IPK2020 | 4,213 | 45,413 |
|  | ZDM01467 | ERS4201447 | IPK2020 | 4,540 | 44,746 |
|  | ZDM02064 | ERS4201446 | IPK2020 | 4,153 | 45,050 |

### Protocol for calling pangene clusters based on WGA evidence

The repository https://github.com/Ensembl/plant-scripts/tree/master/pangenes contains the documentation and source code for calling pangenes. The main script (*get_pangenes.pl*), illustrated in **Figure 1**, sequentially runs the scripts *_cut_sequences.pl*, *_collinear_genes.pl* and *_cluster_analysis.pl*. These tasks can be performed serially on a computer (default) but can also run in batches over a high performance computer cluster. Four types of sequences (cDNA, CDS [amino and nucleotide] and gDNA) are cut so that they can be subsequently added to pangene clusters. cDNA and CDS sequences are cut with GffRead with arguments -w, -y and -x (Pertea & Pertea, 2020). Genomic segments (gDNA) are cut with *bedtools getfasta* (Quinlan & Hall, 2010). WGAs in PAF format are computed with minimap2 with parameters --cs -x asm20 --secondary=no -r1k,5k (Li, 2018) or GSAlign with parameters -sen-no_vcf -fmt 1 (Lin & Hsu, 2020). Unlike minimap2, GSAlign provides ANI estimates. Feature overlap is computed with *bedtools intersect* with parameters -f 0.5 -F 0.5 -e (Quinlan & Hall, 2010) after converting WGAs to BED files, which requires parsing the CIGAR strings contained in PAF files. When features are actual gene models, strandedness is also required. With the exception of dataset ACK2, which includes two genomes with low ANI, the analyses presented in tables 2 and 3 were obtained with *get_pangenes.pl* and optional argument -s, which computes Whole Genome Alignments only with homologous chromosomes. This was indicated with the regular expressions ’^\d+’, ’^\d+[ABD]$’ and ’^chr\d+H’ for rice, wheat and barley respectively. Results can be downloaded at https://github.com/Ensembl/plant-scripts/releases/download/Apr2023/pangenes_bench.tgz (FASTA files not included).

**Figure 1.**
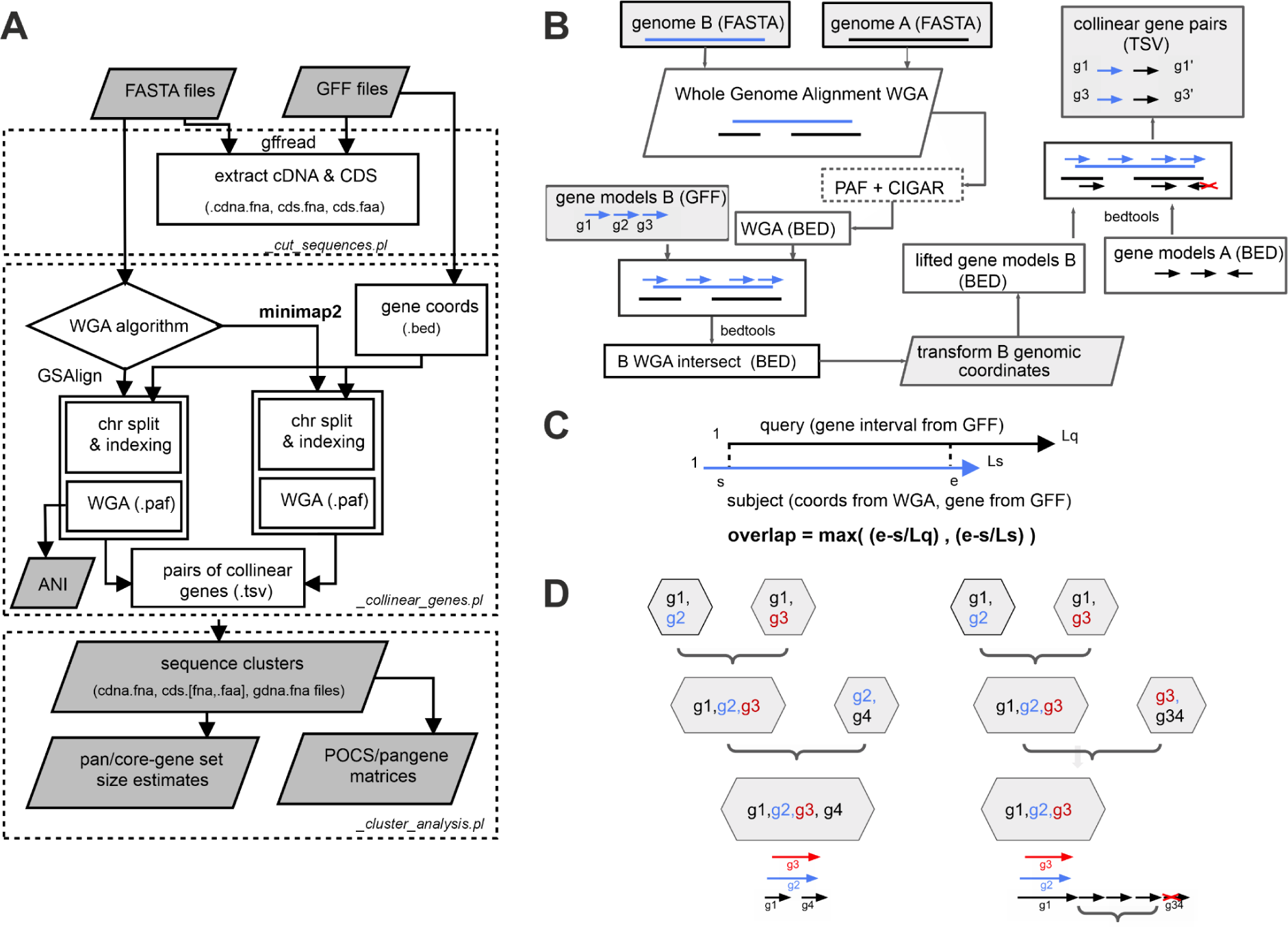
Features of get_pangenes.pl. A) Flowchart of the main tasks and deliverables of script *get_pangenes.pl*: cutting cDNA and CDS sequences (top), calling collinear genes (middle, panels B and C) and clustering (bottom, panel D). By default, only cDNA and CDS sequences longer than 100bp are considered. Whole Genome Alignments (WGA) can be computed with minimap2 (default) or GSAlign, and the input genomes can optionally be split in chromosomes or have their long geneless regions (>1Mbp) masked. Resulting gene clusters contain all isoforms and are post-processed to produce pangene and Percentage of Conserved Sequences (POCS) matrices, as well as to estimate pan-, soft-core-, and core-genomes. GSAlign also produces Average Nucleotide Identity (ANI) matrices. Several tasks can be fine-tuned by customizing an array of parameters, of which alignment coverage is perhaps the most important. B) WGA of genomes A and B produces BED-like files that are intersected with gene models from B. Intersected coordinates are then used to transform B gene models to the genomic space of A. Finally, overlapping A gene models on the same strand are defined as collinear genes. C) Feature overlap is computed from WGAs and gene coordinates from source GFF files. When checking the overlap of A and B gene models, strandedness is required. Overlaps can also be estimated between gene models annotated in one assembly and matched genomic segments from others. D) Making greedy clusters by merging pairs of collinear genes. This algorithm has a key parameter, the maximum distance (in genes) among sequences of the same species that go in a cluster (default=5). Its effect is illustrated on the right side, where gene g34 is left unclustered for having too many intervening genes.

**Table 2.** Summary of pangene analyses based on minimap2 Whole Genome Alignments (WGA). N50 values, that describe the length of aligned genomic fragments, are shown as ranges of observed [min, max] values. The percentage of genes in blocks of 3+ contiguous genes is also shown as a range. Note that barley and wheat datasets require optional argument -H, which masks geneless regions longer than 1Mbp, where repeated sequences accumulate. Maximum RAM use was measured for pairwise WGA batch jobs.

|  | max<br>RAM<br>(GB) | WGA N50<br>(Kbp) | % genes<br>blocks3+ | total<br>clusters | (soft) core<br>clusters | % BUSCO<br>complete |
| --- | --- | --- | --- | --- | --- | --- |
| ACK2 | 4.5 | 6.1 | 34.0 | 38,785 | 20,647 | 94.1 |
| rice3 | 1.4 | [27.4, 29] | [75.1,<br>77.8] | 61,913 | 19,170 | 85.2 |
| chr1wheat10<br>(-H) | 64.5 | [80.8,<br>142.4] | [38.8,<br>53.8] | 25,280 | 9,937 |  |
| barley20 (-H) | 46.3 | [43.6,<br>75.7] | [25.4,<br>35.3] | 165,544 | 23,811<br>(29,226) | 86.6<br>(95.3) |

**Table 3.** Summary of pangene analyses based on GSAlign Whole Genome Alignments (WGA). N50 values, that describe the length of aligned genomic fragments, are shown as ranges of observed [min, max] values. The percentage of genes in blocks of 3+ contiguous genes and the Average Nucleotide Identities (ANI) are also shown as ranges. Maximum RAM use was measured for pairwise WGA batch jobs.

|  | max<br>RAM<br>(GB) | WGA<br>N50<br>(Kbp) | % genes<br>blocks3<br>+ | total<br>clusters | (soft)<br>core<br>clusters | % BUSCO<br>complete | % ANI |
| --- | --- | --- | --- | --- | --- | --- | --- |
| ACK2 | 4.5 | 4.3 | 24.6 | 43,282 | 16,476 | 74.9 | 84.7 |
| rice3 | 3.3 | [15.2,<br>16.9] | [53.0,<br>57.9] | 62,747 | 18,726 | 84.6 | [96.4,<br>97.6] |
| chr1wheat10 | 83.4 | [40.9,<br>72.1] | [20.4,<br>34.3] | 30,005 | 7,833 |  | [98.9,<br>99.4] |
| barley20 | 113.1 | [17.1,<br>34.3] | [10.8,<br>16.2] | 168,880 | 15,567<br>(23,625) | 61.6<br>(82.5) | [96.9,<br>99.3] |

### Pangenome terminology

We will often use pangenome-related terms to describe the pangene clusters output by our protocol. We define occupancy as the number of genomes represented in a cluster. Core clusters contain sequences from all analyzed genomes. Soft-core clusters contain sequences from 95% of the input genomes. Finally, in this paper shell clusters are those with less occupancy than soft-core clusters after excluding singletons (occupancy=1).

### Dotplots

The *_dotplot.pl* script can be used to make a genome-wide dotplot of collinear gene models resulting from a pairwise WGA stored in TSV format. This is done in two steps: i) the TSV file is converted to a PAF file, ii) the dotplot is produced with R package *pafr*, available at https://github.com/dwinter/pafr.

### BUSCO analysis

In order to evaluate the completeness of the core and soft-core collections of pangenes produced by *get_pangenes.pl*, the corresponding protein FASTA files containing all known isoforms of genes were analyzed with BUSCO v5.4.3 (Manni et al., 2021). The poales_odb10 lineage was selected for all datasets except ACK2, where brassicales_odb10 was used instead.

### Venn diagrams

Comparisons of CDS pangene sets produced with both WGA algorithms were carried out with script *compare_clusters.pl* from the GET_HOMOLOGUES-EST software (Contreras-Moreira et al., 2017). As explained in the documentation at https://github.com/Ensembl/plant-scripts/tree/master/pangenes, other scripts from this package can be used to simulate and plot the pangene set growth. The resulting Venn diagrams were plotted with Venn-Diagram-Plotter v1.6.7458 (https://github.com/PNNL-Comp-Mass-Spec/Venn-Diagram-Plotter).

### Ensembl orthologues and InterPro annotations

High-confidence orthogroups produced with Ensembl Compara (Herrero et al., 2016) were retrieved with script *ens_syntelogs.pl* (Contreras-Moreira et al., 2022) from Ensembl Plants (Yates et al., 2022). These are derived from phylogenetic trees of aligned protein sequences from most plant species in Ensembl and have extra supporting collinearity evidence. Only pairs of orthologues with WGA score ≥ 50% and Gene Order Conservation (GOC) ≥ 75% were taken. In other words, only genes with ≥ 50% exonic coverage in Whole Genome Alignments and conserved neighbor gene order were retrieved and considered collinear. Note that GOC allows for inversions and gene insertions.

For ACK2 and rice3 genomes, annotated InterPro protein domains were retrieved from Biomart (Kinsella et al., 2011). These were taken to curate pangene clusters where all their sequences shared at least one InterPro domain.

### Lifting-over gene models on genomic segments

The script *check_evidence.pl* uses precomputed collinearity evidence, stored in a TSV file, and lift-over alignments to project cDNA/CDS sequences on a reference genomic sequence. Briefly, collinear genome sequences are extracted with *bedtools getfasta* and then pre-clustered cDNA/CDS sequences are mapped to them with GMAP, a software tool that efficiently connects exons while accurately defining splice sites and jumping intervening introns (Wu & Watanabe, 2005). GMAP is run with parameters -t 1 -2 -z sense_force -n 1 -F. Increasing the verbosity of the script produces the actual GMAP sequence alignments, which are useful to inspect failed lift-over attempts (ie partially aligned proteins, premature stop codons, length not multiple of 3).

### Flowering genes

A collection of 26 genes relevant in barley breeding due to their roles in flowering control and spike architecture was compiled. Pangene clusters were produced for the union of the barley20 dataset and two more high-quality gene annotations: MorexV3 (Mascher et al., 2021) and BaRTv2 (Coulter et al., 2022). The former is the IPK annotation from http://doi.org/10.5447/ipk/2021/3 (35,826 gene models, assembly GCA_904849725.1) and the latter the JHI annotation from https://ics.hutton.ac.uk/barleyrtd/bart_v2_18.html (39,281 gene models, assembly ERS4201450). The script *get_pangenes.pl* was run with arguments -s ’^chr\d+H’ -H -t 0. The resulting clusters were aligned with Clustalx 2.1 for manual curation (Larkin et al., 2007).

## RESULTS AND DISCUSSION

### A protocol for calling pangenes based on Whole Genome Alignments

The first result from this work is the design of a protocol for calling pangenes in a series of related genomes. A pangene is defined as a gene model found within a homologous region in a set of genomes. In order to find pangenes, WGAs are computed, which in turn produce pairs of collinear genomic segments. Collinear evidence is stored in TSV files and can be produced by two WGA algorithms: minimap2 and GSAlign. The protocol is represented as a flowchart in **Figure 1A**. As illustrated schematically in **Figure 1B**, WGAs are used to project the coordinates of gene models across assemblies. By default, a pair of genes are said to be collinear when at least half the length of one matches the other in genomic space (**Figure 1C**). Finally, clusters of genes (pangenes) emerge by merging pairs of collinear genes from different input taxa (**Figure 1D**).

An example collinear region of *Oryza sativa* Japonica group (bottom) and *Oryza nivara* (dataset rice3) as displayed in the Ensembl Plants genome browser is shown in **Figure 2**, together with a summary of the supporting WGA evidence. Besides five 1-to-1 collinear gene pairs, it can be seen that the gene ONIVA01G00100 was mapped to two consecutive *O. sativa* models (Os01g0100100, Os01g0100200) and that two *O.sativa* genes map to unannotated genome segments in *O. nivara*. In a nutshell, this figure shows that WGA evidence allows matching of long genes to split genes if they are collinear, as well as matching annotated gene models to homologous regions in other genomes, even if the genes in question failed to be annotated.

**Figure 2.**
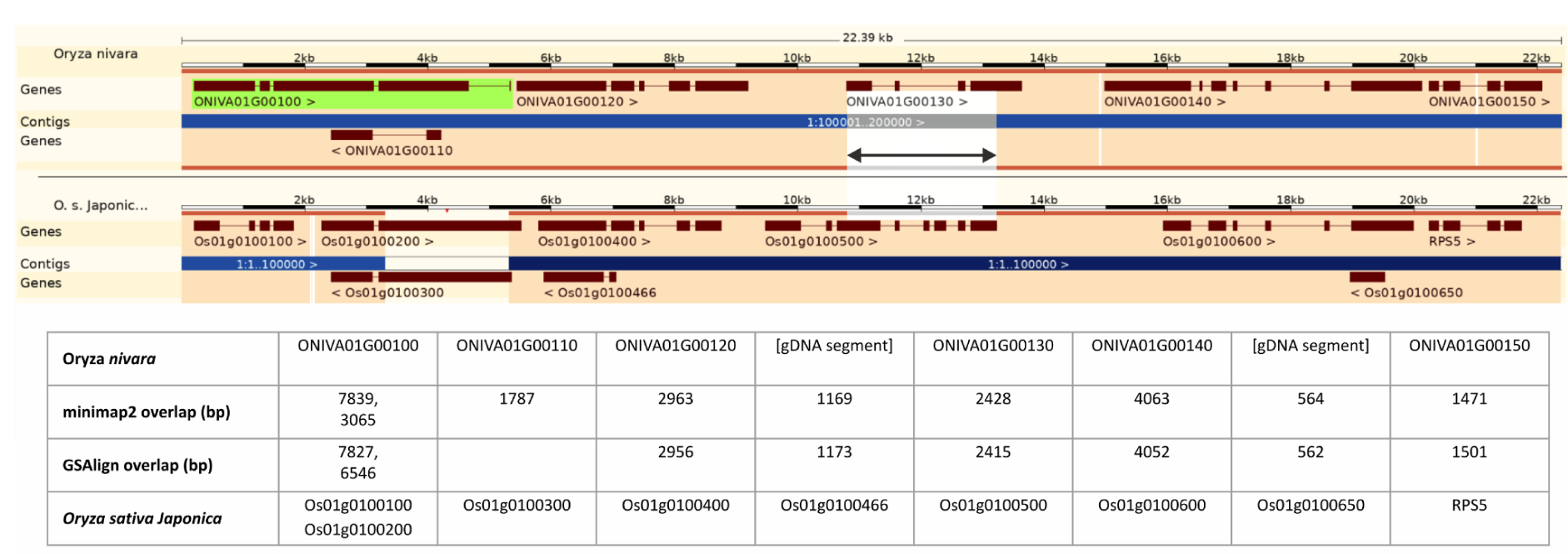
Aligned genomic region in chr1 of *Oryza nivara* (top) and *Oryza sativa* Japonica group cv. Nipponbare (bottom) as displayed in the Ensembl Plants browser. Genes on the forward strand (>) are above contigs, whilst those in the negative strand (<) are underneath. As a result of the genomic alignment, genes of *O. nivara* overlap with gene models from *O. sativa*. This evidence can be used to identify collinear genes that take equivalent positions in different genomes, as illustrated with gene models ONIVA01G00130 and Os01g0100500, which overlap over 2.4kb (white rectangle). The example shows that overlapping gene models might share only some exons. The table below shows the collinear gene models identified based on minimap2 ang GSAlign alignments, together with the corresponding overlapped based pairs.

### Benchmark on several plant datasets

A systematic benchmark of *get_pangenes.pl* was performed with the four datasets in **Table 1**. In order to describe the performance several variables were collected. As computing WGAs is costly, particularly for large genomes, the maximum amount of RAM consumed by pairwise genome alignments was recorded. The size of the collinear fragments produced by WGA is captured in two variables, N50 and the percentage of fragments that contain blocks of 3 or more genes. Several further variables were also calculated to describe the clusters of collinear genes produced by the protocol: the total number of clusters, the number of pangenes present in all genotypes (core clusters), the pangenes present in 95% of genotypes (soft-core clusters) and the percentage of complete BUSCOs, which are universal single-copy orthologs. The results are summarized on **Table 2** (minimap2) and **Table 3** (GSAlign). Here the outcomes of both algorithms are compared. Although GSAlign consumes more RAM than minimap2 as genome size grows, this is not a fair comparison; in fact, while GSAlign was fed raw genome sequences, minimap2 only completed the WGAs for barley and wheat after masking long geneless genomic regions. The collinear segments aligned by minimap2 are longer and contain more genes than those produced by GSAlign. **Supplemental Figures S1 and S2** confirm that collinear gene models have large overlaps and that the genomic segments that contain them are generally syntenic along chromosomes. **Supplemental Table S1** shows that the number of hits per gene is similar for both algorithms, with minimap2 failing to map more genes than GSAlign. Despite these differences, the numbers of pangenes clustered using WGA evidence from both algorithms are comparable: 86.3% of all clusters are identical (95.5% for core clusters, see **Supplemental Figure S3**).

As an extra test, clusters of cDNA and CDS sequences resulting from the rice3 analysis were aligned locally to compute their sequence identity both at the nucleotide and protein level. Note that these clusters contain all isoforms annotated, so often there might be several sequences for the same gene. The results, plotted in **Supplemental Figure S4**, yielded median sequence identities of 99.6% for nucleotides (cDNA and CDS). For protein sequences the median values are 98.3% (GSAlign) and 98.1 % (minimap2). This means that annotated sequences clustered together are nearly identical, although, as seen in **Supplemental Figure S5**, that does not guarantee that the same protein sequence is always encoded by clustered genes, as a result of divergent gene model annotation. Note that we also found some cases, 273/30,705 for minimap2 and 320/30,129 for GSAlign, where cDNA sequences of the same cluster could not be aligned. These occur when overlapping genes do not share exons. Nevertheless, these pangenes are not filtered out by default as such gene models could encode loss of function alleles that might be valuable to capture.

**Tables 2** and **Table 3** also contain a summary of BUSCO analysis. BUSCOs are sets of universal single-copy orthologs tailored to different taxa and are typically used to estimate the completeness of genome assemblies. In this context, BUSCOs provide a biologically meaningful metric based on expected gene content (Manni et al., 2021). The ACK2 dataset contains the two most divergent genomes, with 84.7% nucleotide identity (see **Table 3**). The core pangenes found across *A. thaliana* and *A. lyrata* contain a higher percentage of complete BUSCOs with minimap2 than with GSAlign (94.1% vs 74.9%). In the rice3 dataset the nucleotide identity rises to 96% and the core pangene set contains ca. 85% complete BUSCOs with both WGA algorithms. As BUSCO analysis does not make sense for a single chromosome, the wheat dataset was left out. As for the barley20 dataset, the nucleotide identity is generally higher than that of rice3 and the benchmark produced core sets with close 86.6% (minimap2) and 61.6 % (GSAlign) complete BUSCOs. This number increased to 95.3% and 82.5% when all soft-core pangenes are considered, revealing a superior performance of minimap2 in this dataset. To put all these BUSCO scores in perspective please see the scores of individual input genome annotations in **Supplemental Table S2**. Finally, **Table 3** also shows the Average Nucleotide Identity values computed by the GSAlign algorithm for pairwise genome alignments. These values are useful to measure the divergence of the genomes being compared.

### Comparison to ancestral karyotype and Ensembl orthogroups

Additional analyses were carried out to gain insights into the performance of our protocol by comparing our results to data produced independently. First, we estimated its recall on the ACK2 dataset, which represents the most difficult scenario tested due to having the lowest nucleotide identity. This experiment counted the number of collinear genes identified by minimap2 and GSAlign within 23 blocks of the Ancestral Crucifer Karyotype. The results, summarized in **Supplemental Table S3**, indicate that in these conditions 65% (minimap2) and 52% (GSAlign) of the genes making up the blocks are called collinear.

Second, taking advantage of the fact that the genomes in datasets ACK2 and rice3 are included in Ensembl Plants, it was possible to compare the pangenes to precomputed Ensembl Compara orthogroups. In this comparison, clusters are said to match orthogroups when they include all the orthologues annotated in Ensembl; the results are shown in **Table 4**. The analysis of the challenging ACK2 dataset shows that *get_pangenes.pl* recovers 90.3% (minimap2) and 70.2% (GSAlign) of core clusters found among Compara orthogroups, while producing more clusters with multiple copies. This means that ANI limits the recall of our protocol more severely for GSAlign than for minimap2. Nevertheless, 91% (18,792/20,647 minimap2) and 89.9% (14,817/16,476 GSAlign) of pangenes group together proteins that share InterPro domains, similar to Compara orthogroups (18,259/20,174; 90.5%). This indicates that pangene clusters are generally biologically relevant.

**Table 4.** Summary of pangene clusters obtained for datasets ACK2 and rice3 and the corresponding orthogroups in Ensembl Plants. Core clusters contain genes from all analyzed genomes; in rice, shell clusters contain genes from two species. Clusters with multiple copies have several genes from the same species. gDNA segments are shell clusters that bring together a gene model and a matching genomic segment from the underlying WGA. Column ‘match Compara’ shows the number of pangene clusters that contain the same genes as the corresponding Compara orthogroups. The last column shows the number of pangene clusters that contain sequences that share an InterPro domain (the number in square brackets is for core clusters only).

|  | dataset | core clusters | multiple copies | shell clusters | gDNA segments | match Compara | share InterPro domains |
| --- | --- | --- | --- | --- | --- | --- | --- |
| Compara orthogroups | ACK2 | 20,192 | 161 |  |  |  | [18,259] |
| minimap2 clusters | ACK2 | 20,647 | 731 |  |  | 18,245 | [18,792] |
| GSAlign clusters | ACK2 | 16,476 | 454 |  |  | 14,181 | [14,817] |
| Compara orthogroups | rice3 | 13,020 | 219 | 6,386 |  |  | 16,766<br>[11,571] |
| minimap2 clusters | rice3 | 22,880 | 3,360 | 7,825 | 6,521 | 18,281 | 23,062<br>[19,239] |
| GSAlign clusters | rice3 | 20,399 | 2,885 | 9,730 | 6,103 | 17,103 | 22,834<br>[17,135] |

The rice3 benchmark revealed that 79.9% (minimap2) and 83.8 (GSAlign) % of pangene clusters match Compara orthogroups, with *get_pangenes.pl* calling over seven thousand more core pangenes than Compara. As a quality check of these core pangenes, we counted how many encoded proteins that share at least one InterPro domain. We found that 84.1% (19,239/22,880 minimap3) and 84% (17,135/20,399 GSAlign) of clusters are consistent in functional terms, compared to 89% (11,571/12,997 Compara). Inspection of the examples on **Supplemental Figure S6**, demonstrate that our protocol is able to cluster together overlapping gene models which might be split or incomplete. This would illustrate why pangene clusters are more likely to group together multiple sequences from the same species. Note that incomplete gene models, or clusters with sequences of contrasting length, would in turn explain why some clusters contain sequences that do not encode a common protein domain (see **Supplemental Figure S7**). The table also shows that over six thousand shell pangene clusters are produced that pair an annotated gene model with an overlapping genomic segment (gDNA) from a different species. **Supplemental Table S4** shows how core and shell clusters are represented as a BED-like pangene matrix produced by *get_pangenes.pl*.

### Confirming gene Presence-Absence Variation

A use case of pangenome analysis of plants and other organisms is to find genes which might be present and functional only in some individuals or populations. These would be annotated by our pipeline as shell genes. As discussed in the previous section, the *get_pangenes.pl* protocol can produce gDNA sequence clusters that contain the genomic intervals of annotated gene models plus overlapping unannotated segments from other species. This feature takes advantage of pre-computed WGAs, which match genomic segments whether they harbor gene models or not. Such clusters can effectively be used to lift-over or project gene models from the species where they are annotated to other individuals. In particular, CDS or cDNA isoform sequences from annotated genes can be mapped to the matching genomic segments and the resulting alignment will directly confirm whether exon/intron boundaries and the embedded coding sequence are conserved. When conserved, the segment likely contains an overlooked gene model; when not, it probably contains a gene fragment or a pseudogene. A flowchart in **Supplemental Figure S7** summarizes how the script *check_evidence.pl* retrieves the WGA evidence supporting a pangene cluster, defines consensus and outlier isoform sequences, and then projects consensus CDS or cDNA sequences on candidate genomic segments with GMAP (See Materials and Methods). Note that, in addition to missing gene models, lifting-over can also merge split gene models or, conversely, divide gene models that might have been merged during gene annotation. Either way, when the projection succeeds and a complete open reading frame is aligned, a patch GFF file is created that conveys the genomic coordinates of the projected gene.

Using shell CDS clusters of occupancy >9 resulting from the minimap2 analysis of dataset barley20 we carried out a survey to see how often the different scenarios (missing, split, merged gene) occur in a real dataset. Out of 41,655 clusters, our approach detected 74 cases where a long model could be potentially corrected, 30 cases of incorrectly split genes and 9,839 potentially missing genes. We selected one candidate missing gene to illustrate the most common situation, pangene Horvu_MOREX_1H01G011400. In this example the original pangene grouped together gene models from 13 barley genotypes, supported by the WGA evidence shown in **Supplemental Table 5**. When CDS nucleotide sequences from those 13 cultivars were aligned to candidate genomic segments of the remaining genotypes, a perfect match was found in the genome sequence of OUN333. The encoded lifted-over protein sequence is shown at the bottom of the multiple alignment in **Figure 3**, being identical to others in the cluster. The resulting patch GFF file that would add this gene model to OUN333 is shown in **Figure 3B**. This test case suggests the combination of WGA evidence and gene model projection could be a powerful way to refine gene annotation across individuals of the same species. Moreover, as shown in the example, lift-over alignments can be used to confirm or reject observed PAV. In this case we can hypothesize this gene model is actually present in cultivar OUN333, and probably missing in the remaining genotypes. Additional data beyond sequence evidence would be required to fully characterize such genes, such as expression data and, ultimately, proteomics evidence.

**Figure 3.**
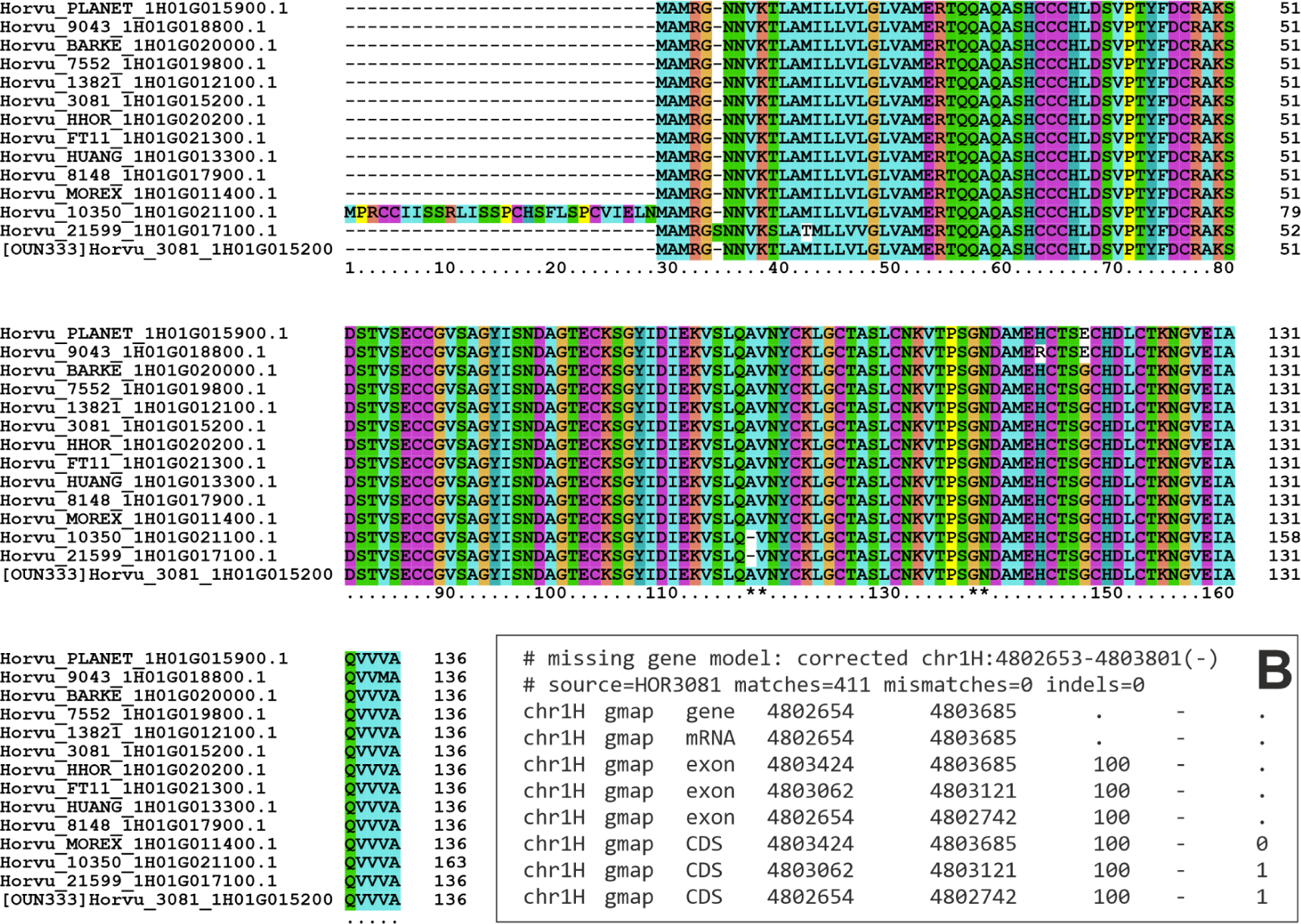
Clustalx multiple alignment of protein sequences encoded in barley pangene cluster Horvu_MOREX_1H01G011400. This cluster contains isoforms from 13 gene models, but none from genotype OUN333. The last sequence is encoded by a CDS sequence lifted-over from cultivar HOR3081 on the genome of OUN3, spanning 3 exons (exon boundaries are marked with asterisks. B) Patch GFF file with the coordinates of the exons lifted-over from gene model Horvu_3081_1H01G015200. The underlying CDS nucleotide sequence was aligned with 411 matches, no indels and no mismatches with check_evidence.pl -f.

### Curation of barley flowering genes

The protocol for pangene clustering was further tested with an increased set of barley assemblies and gene annotations. In this final experiment the barley20 dataset plus the high-quality MorexV3 and BaRTv2 gene annotations were pooled and a subset of 26 genes known to regulate flowering and spike architecture extracted from the resulting clusters. The curated results are summarized on **Table 5**, where it can be seen that in all cases these genes were found in 20 or more barley annotations. The results in this gene survey were curated and we found some cases (HvFT3/Ppd-H2, HvLUX, HvGRP7b, HvLHY) where a gene was missing from a cluster in some cultivars but there was a candidate genomic region harboring part of that sequence. That would be the case for cultivars Igri and HOR3081 and locus HvFT3/Ppd-H2. As explained in the previous section, we lifted-over the collinear CDS nucleotide sequences from all the other cultivars and we obtained identical matches but only for exon 4 (see **Supplemental Figure S9**), replicating previous observations that exons 1 to 3 of this gene have been deleted in some genotypes (Kikuchi et al., 2009).

**Table 5.**
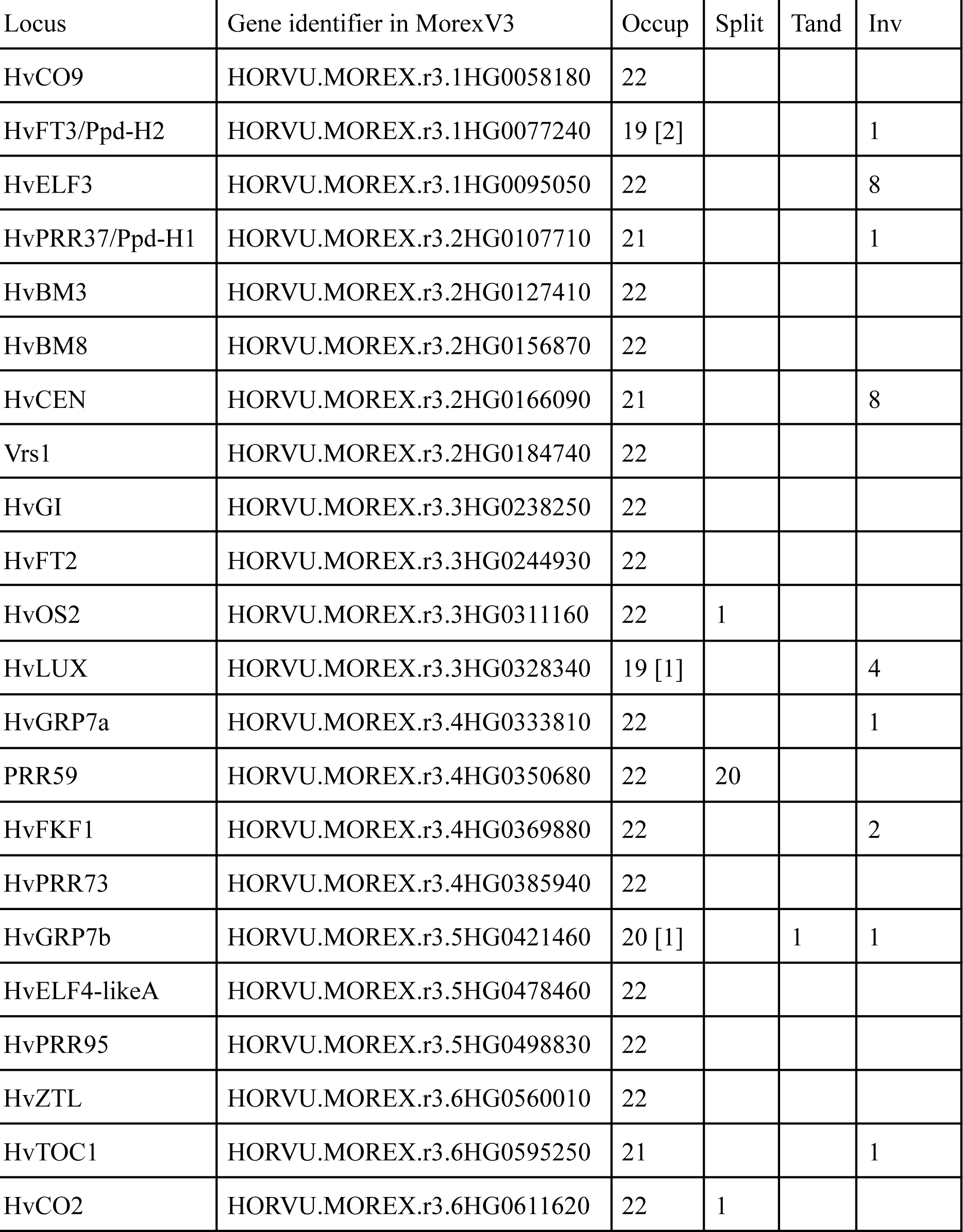
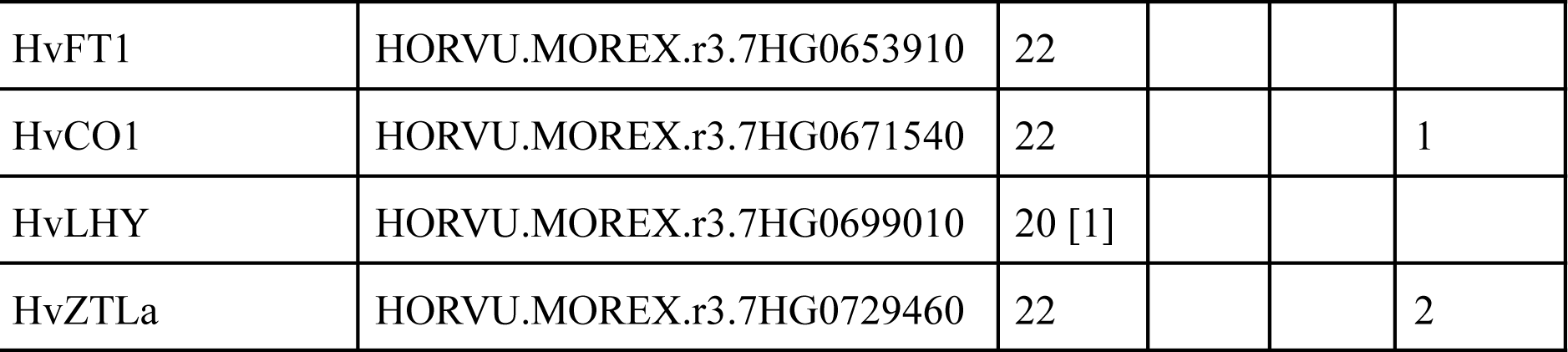
Survey of flowering-related pangenes in a collection of 22 barley genotypes, including the barley20 dataset plus the MorexV3 and BaRTv2 high-quality annotations. Column ‘Occup’ indicates how many annotations contain a gene model, with the number in brackets being the number of matched genomic segments in cases where a gene was absent in some cultivars. Column ‘Split’ tells how many gene models are split in each pangene cluster with respect to the mode gene model. Column ‘Tand’ says how many extra tandem gene models are in each cluster. Column ‘Inv’ states how many genes were found inverted in Whole Genome Alignments.

Another interesting case was locus HvPRR37/Ppd-H1, which was found to be absent also in cultivar Igri, despite the gene being cloned in this cultivar with accession AY970701.1 (Turner et al., 2005). As it turns out, the Igri genome assembly placed this gene model (Horvu_IGRI_Un01G026500) outside of the pseudo-chromosomes; instead it is located in chrUn, and therefore it cannot be collinear to the homologous genes in other cultivars when only homologous chromosomes are compared (option -s).

We also found several instances where genes were found in inverted genome regions, known to be valuable to reconstruct the history of crops (Zhou et al., 2023). In the case of HvCEN this observation matches previous reports (Jayakodi et al., 2020), but we found other cases such as HvELF3 or HvLUX. These examples show the value of using Whole Genome Alignments for the definition of pangenes, as the strand of genes conveys chromosomal and evolutionary information.

Finally, in this set of genes used by barley breeders we also observed instances of gene models found to be split in some annotations (see **Supplemental Figure S10**) or genes with extra tandem copies. An example of the former is cluster HORVU.MOREX.r3.3HG0311160, which corresponds to barley locus HvOS2, which encodes MADS-box protein ODDSOC2 (Greenup et al., 2010), is depicted in **Figure 4**.

**Figure 4.**
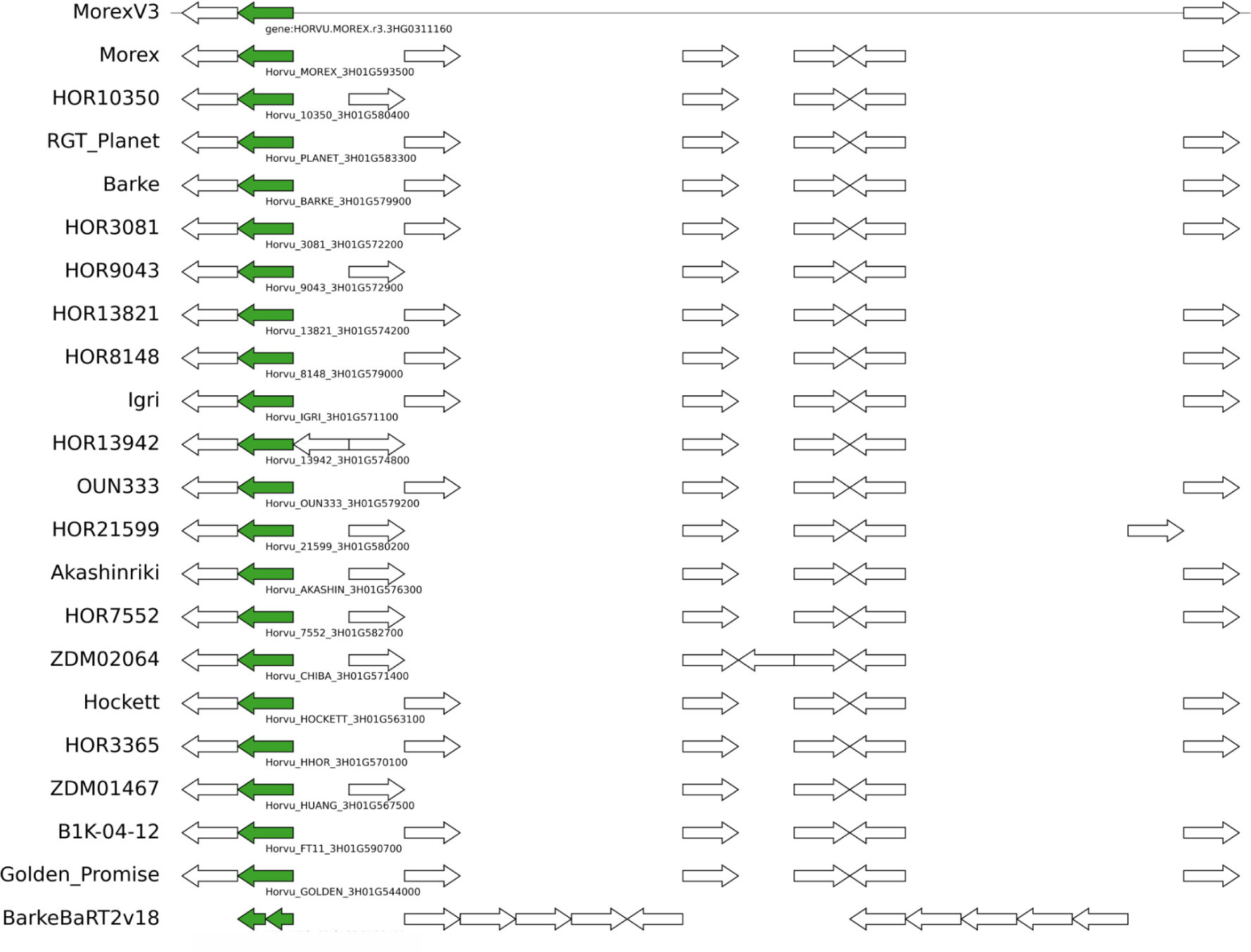
Genomic context of pangene cluster HORVU.MOREX.r3.3HG0311160 (green arrows), which corresponds to barley locus HvOS2. The genome fragment on top corresponds to reference genome MorexV3 and the tracks below show collinear genes found in other barley assemblies and annotation sets. In this example the BarkeBaRT2v18 gene is split in two partial models. Note that white gene models might not be collinear as they could be encoded in a different genome fragment. Figure generated with script check_evidence.pl and pyGenomeViz (https://github.com/moshi4/pyGenomeViz).

Altogether, these results highlight the challenges of consistently annotating gene models in individuals of the same species and confirm that analyzing soft-core, instead of core genes, is probably a good idea to tolerate the situations described in this section.

## ACKNOWLEDGEMENTS

We are grateful to all members of the PanOryza consortium for feedback during the development of this work. We thank the Gramene team for continuous support and cooperation, as well as members of the Ensembl team for developing and maintaining the front-end and back-end software and infrastructure that underpins Ensembl Plants [Wellcome Trust WT222155/Z/20/Z]. We would like to thank BBSRC/NSF for funding [BB/T015691/1, BB/T015608/1], CSIC for funding [FAS2022_052] and the European Molecular Biology Laboratory.

## CONFLICT OF INTEREST

Paul Flicek is a member of the Scientific Advisory Boards of Fabric Genomics, Inc. and Eagle Genomics, Ltd.

## ORCID

0000-0002-5462-907X Bruno Contreras Moreira 0000-0002-6433-8356 Shradha Saraf

0000-0002-0523-4071 Guy Naamati

0000-0003-3484-2655 Ana M Casas

0000-0002-8965-1648 Sandeep S Amberkar 0000-0002-3897-7955 Paul Flicek

0000-0001-6118-9327 Andrew R Jones

0000-0001-5690-9633 Sarah Dyer

## SUPPLEMENTAL MATERIAL

Supplementary tables and figures are included in file Supplementary_data.pdf.

## Notes

https://github.com/Ensembl/plant-scripts/releases/download/Apr2023/pangenes_bench.tgz

### Summary of Updates

* updated algorithm now handles by default inverted genomic segments * added benchmark on curated flowering-related genes in barley * new figures * new supplementary data added

