## Supplementary tables and figures for "Calling pangenes from plant genome alignments confirms presence-absence variation"

**Table S1**. Other Whole Genome Alignment stats for minimap2 and GSAlign algorithms. The different variables are shown as ranges of [min, max] observed values among all pairwise alignments in a dataset.

| | minimap2 | | GSAlign | |
|---|---|---|---|---|
| dataset | hits/gene | unmapped genes | hits/gene | unmapped genes |
| rice3 | [1.020, 1.117] | [140, 632] | [1.019, 1.116] | [6, 177] |
| chr1wheat10 | [1.021, 1.033] | [282, 513] | [1.005, 1.011] | [13, 200] |
| barley20 | [1.011, 1.050] | [55, 3259] | [1.004, 1.013] | [0, 741] |

**Table S2**. Summary of BUSCO completeness analyses of individual genomes that are part of datasets in this paper. BUSCO percentages are shown as ranges of observed [min, max] values among annotated genomes in a dataset.

| | lineage dataset | % BUSCO complete |
|---|---|---|
| ACK2 | brassicales_odb10 | [95.9, 100] |
| rice3 | poales_odb10 | [84.7, 95.7] |
| barley20 | poales_odb10 | [97.7, 98.6] |

**Table S3**. Collinear genes found between *Arabidopsis thaliana* and *A. lyrata* within 23 blocks of the Ancestral Crucifer Karyotype (Lysak et al., 2016) based on Whole Genome Alignments produced with minimap2 and GSAlign. Blocks are defined as lists of contiguous genes in *A. thaliana*.

| block | boundary genes | #genes | minimap2 | GSAlign |
|---:|---:|---:|---:|---:|
| A | [AT1G01010,AT1G19840] | 2388 | 1724 | 1620 |
| B | [AT1G19850,AT1G37130] | 1932 | 1177 | 976 |
| C | [AT1G43020,AT1G56190] | 1314 | 789 | 817 |
| D | [AT1G56210,AT1G64670] | 945 | 523 | 110 |
| E | [AT1G64960,AT1G80950] | 1993 | 1377 | 1279 |
| F | [AT3G01015,AT3G25520] | 3118 | 2162 | 2083 |
| G | [AT2G05170,AT2G07690] | 274 | 75 | 76 |
| H | [AT2G10940,AT2G20900] | 903 | 519 | 412 |
| I | [AT2G20920,AT2G31035] | 1240 | 818 | 766 |
| J | [AT2G31040,AT2G48150] | 2218 | 1514 | 1571 |
| KL1 | [AT2G01060,AT2G05160] | 479 | 272 | 279 |
| KL2 | [AT3G25540,AT3G32960] | 693 | 359 | 368 |
| MN | [AT3G42180,AT3G63530] | 2595 | 1700 | 1735 |
| O | [AT4G00026,AT4G05450] | 705 | 401 | 112 |
| P | [AT4G07390,AT4G12620] | 609 | 314 | 322 |
| Q | [AT5G23010,AT5G30510] | 720 | 430 | 366 |
| R | [AT5G01010,AT5G23000] | 2541 | 1864 | 1786 |
| S | [AT5G32470,AT5G42110] | 881 | 459 | 154 |
| T | [AT4G12700,AT4G16240] | 522 | 335 | 22 |
| U | [AT4G16250,AT4G40100] | 3039 | 2137 | 274 |
| V | [AT5G42130,AT5G47810] | 727 | 459 | 28 |
| W | [AT5G47820,AT5G60800] | 1613 | 1103 | 1123 |

| | | | | |
|---|---|---|---|---|
| X | [AT5G60805,AT5G67640] | 865 | 631 | 479 |
| total | | 32314 | 21142 | 16758 |

**Table S4**. Excerpt from BED-like pangene matrix produced during the analysis of dataset rice3. Note that non-reference/unplaced genes appear as comments (#) but placed in their likely pan-genomic location according to their position in non-reference genomes. In this example the reference is the genome of *Oryza sativa* Japonica Group cv. Nipponbare. 'Occup' stands for pangenome occupancy, the number of genomes where a pangene was found. By default pangenes take their names from individual genes, but they could be renamed with any type of identifiers.

| chr | start | end | pangene | occup | strand | *O. sativa* Japonica | *O. nivara* | *O. sativa* Indica |
|---|---|---|---|---|---|---|---|---|
| #1 | NA | NA | ONIVA01G00090 | 1 | 0 | NA | ONIVA01G00090 | NA |
| 1 | 2983 | 10815 | Os01g0100100 | 3 | + | Os01g0100100 | ONIVA01G00100 | BGIOSGA002569 |
| 1 | 11218 | 12435 | Os01g0100200 | 2 | + | Os01g0100200 | NA | BGIOSGA002570 |
| 1 | 11372 | 12284 | Os01g0100300 | 3 | - | Os01g0100300 | ONIVA01G00110 | BGIOSGA002567 |
| 1 | 12721 | 15685 | Os01g0100400 | 3 | + | Os01g0100400 | ONIVA01G00120 | BGIOSGA002571 |
| 1 | 12808 | 13978 | Os01g0100466 | 1 | - | Os01g0100466 | NA | NA |
| 1 | 16399 | 20144 | Os01g0100500 | 3 | + | Os01g0100500 | ONIVA01G00130 | BGIOSGA002572 |
| 1 | 22841 | 26892 | Os01g0100600 | 3 | + | Os01g0100600 | ONIVA01G00140 | BGIOSGA002573 |
| 1 | 25861 | 26424 | Os01g0100650 | 1 | - | Os01g0100650 | NA | NA |
| 1 | 27143 | 28644 | Os01g0100700 | 3 | + | Os01g0100700 | ONIVA01G00150 | BGIOSGA002574 |
| 1 | 29818 | 34453 | Os01g0100800 | 3 | + | Os01g0100800 | ONIVA01G00160 | BGIOSGA002575 |
| 1 | 35623 | 41136 | Os01g0100900 | 3 | + | Os01g0100900 | ONIVA01G00170 | BGIOSGA002576 |
| #1 | NA | NA | BGIOSGA002577 | 1 | 0 | NA | NA | BGIOSGA002577 |
| 1 | 58658 | 61090 | Os01g0101150 | 2 | + | Os01g0101150 | NA | BGIOSGA002578 |

**Table S5**. Summary of Whole Genome Alignment (WGA) evidence for the gene models in CDS cluster Horvu_MOREX_1H01G011400 resulting from the analysis of dataset barley20. This cluster contains isoforms from 13 gene models. Note that there is no gene from barley genotype OUN333. Column 'pairs' indicates how many WGA alignments relate a gene model to other models in the same cluster, with column 'overlap' summing up all overlapping genomic regions in those WGAs.

| length | pairs | overlap | gene name | taxon |
|---:|---:|---:|---|---|
| 1118 | 12 | 11848 | Horvu_10350_1H01G021100 | HOR10350 |
| 1034 | 11 | 11024 | Horvu_BARKE_1H01G020000 | Barke |
| 1036 | 11 | 11266 | Horvu_21599_1H01G017100 | HOR21599 |
| 1034 | 11 | 9737 | Horvu_HUANG_1H01G013300 | ZDM01467 |
| 1034 | 10 | 10029 | Horvu_HHOR_1H01G020200 | HOR3365 |
| 1034 | 10 | 10054 | Horvu_FT11_1H01G021300 | B1K-04-12 |
| 1031 | 10 | 10203 | Horvu_MOREX_1H01G011400 | Morex |
| 1031 | 10 | 10199 | Horvu_3081_1H01G015200 | HOR3081 |
| 1033 | 10 | 10065 | Horvu_8148_1H01G017900 | HOR8148 |
| 1029 | 9 | 9280 | Horvu_9043_1H01G018800 | HOR9043 |
| 1034 | 8 | 8114 | Horvu_13821_1H01G012100 | HOR13821 |
| 1028 | 6 | 6188 | Horvu_PLANET_1H01G015900 | RGT_Planet |
| 1033 | 4 | 3823 | Horvu_7552_1H01G019800 | HOR7552 |

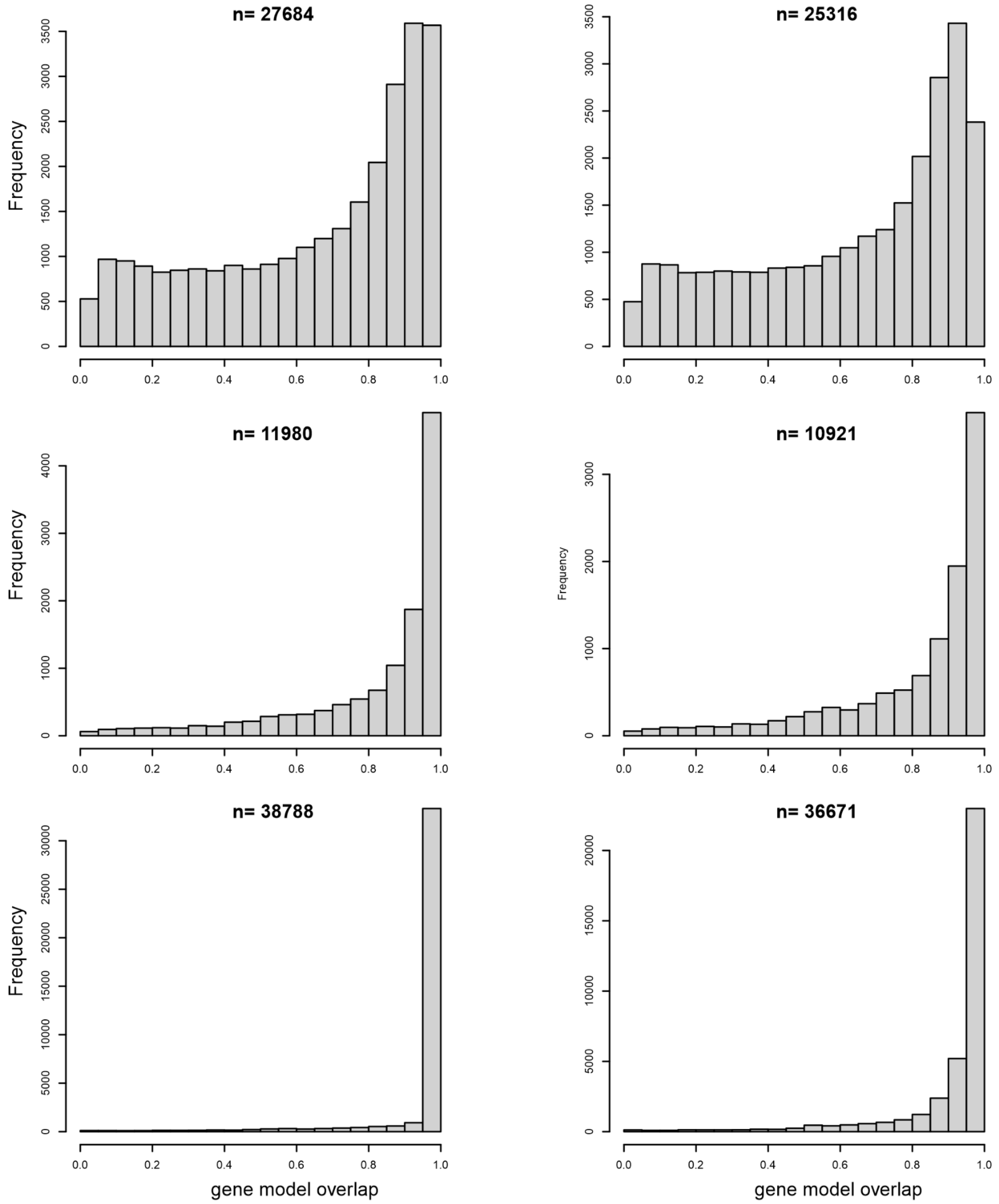


**Figure S1.** Overlap ratio of collinear gene models in rice, wheat and barley. Results based on minimap2 Whole Genome Alignments are on the left, GSAlign results on the right. Top) *Oryza nivara* and *Oryza sativa* Japonica group (rice3 dataset). Middle) Chinese Spring and Julius (chr1wheat10 dataset). The minimap2 analysis was carried out with optional parameter -H, which masks geneless regions longer than 1Mbp, where repeated sequences accumulate.

Bottom) Morex and Barke (barley20 dataset), with the minimap2 analysis done with parameter -H. Ratios in the plots are calculated with respect to coordinates in the source GFF files. However, overlap in *_collinear_genes.pl* is computed with respect to WGA alignments, which might be partial. That explains why there are cases with the default overlap ratio < 0.5.

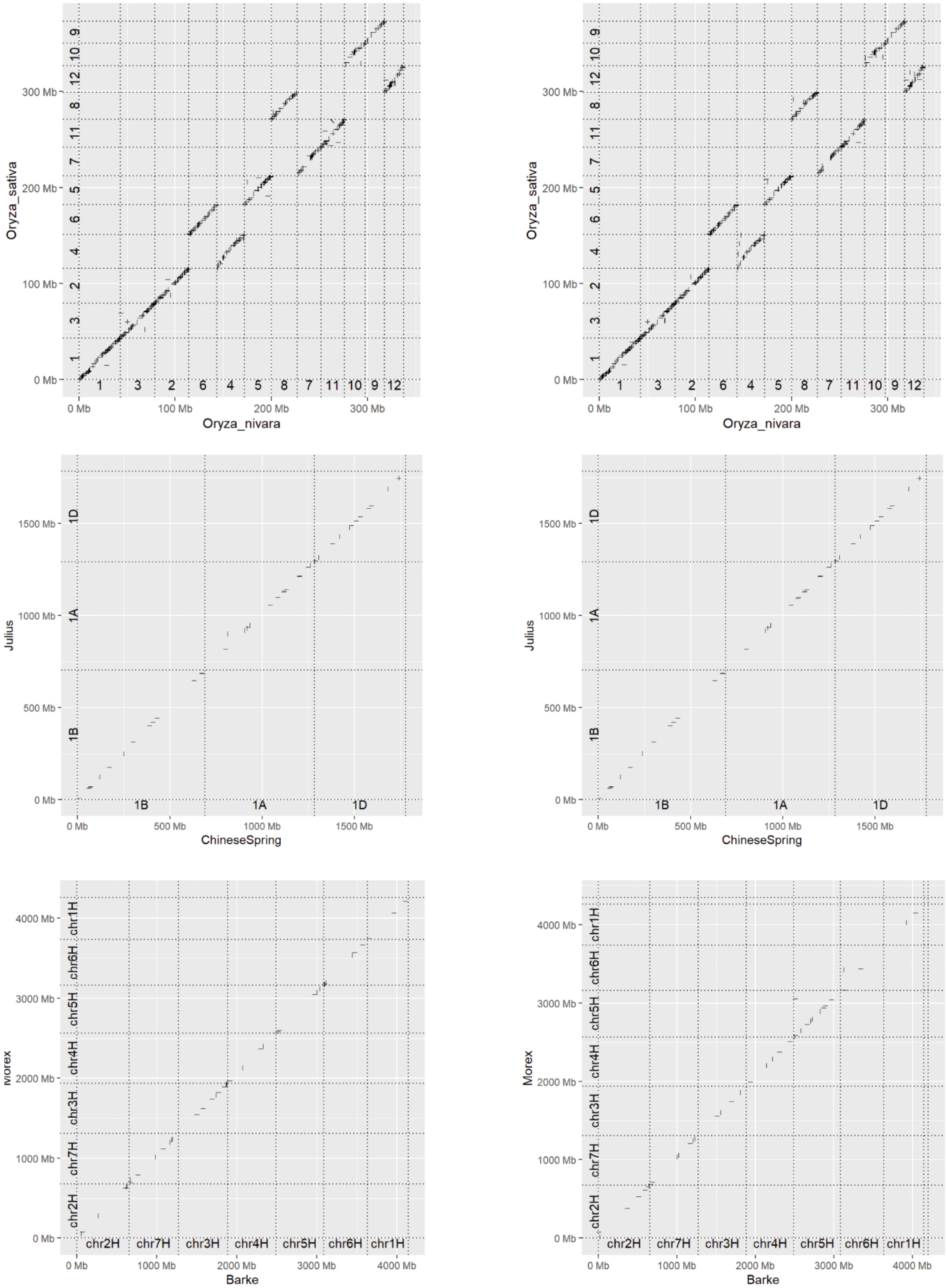


**Figure S2.** Dot plots of collinear gene models called in rice, wheat and barley genomes. Note that chromosomes are sorted by size. Results based on minimap2 Whole Genome Alignments are shown on the left, with GSAlign-based results on the right. A) *Oryza nivara* and *Oryza sativa* Japonica group (rice3 dataset). B) Chinese Spring and Julius (chr1wheat10 dataset). The minimap2 analysis was carried out with optional parameter -H, which masks geneless regions longer than 1Mbp, where repeated sequences accumulate. C) Morex and Barke (barley20 dataset), with the minimap2 analysis done with parameter -H. Plots created with R package *pafr*.

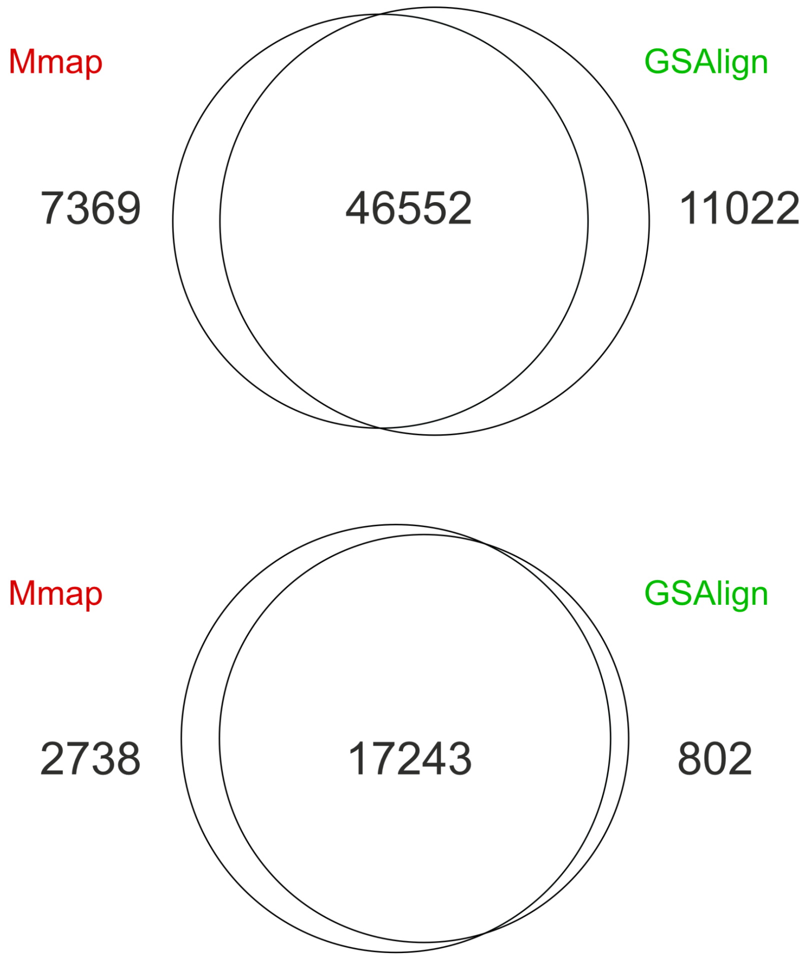


**Figure S3.** Venn diagrams of pangene clusters based on minimap2 and GSAlign Whole Genome Alignments of the rice3 dataset. Top) CDS nucleotide clusters of all occupancies. Bottom) Core CDS nucleotide clusters.

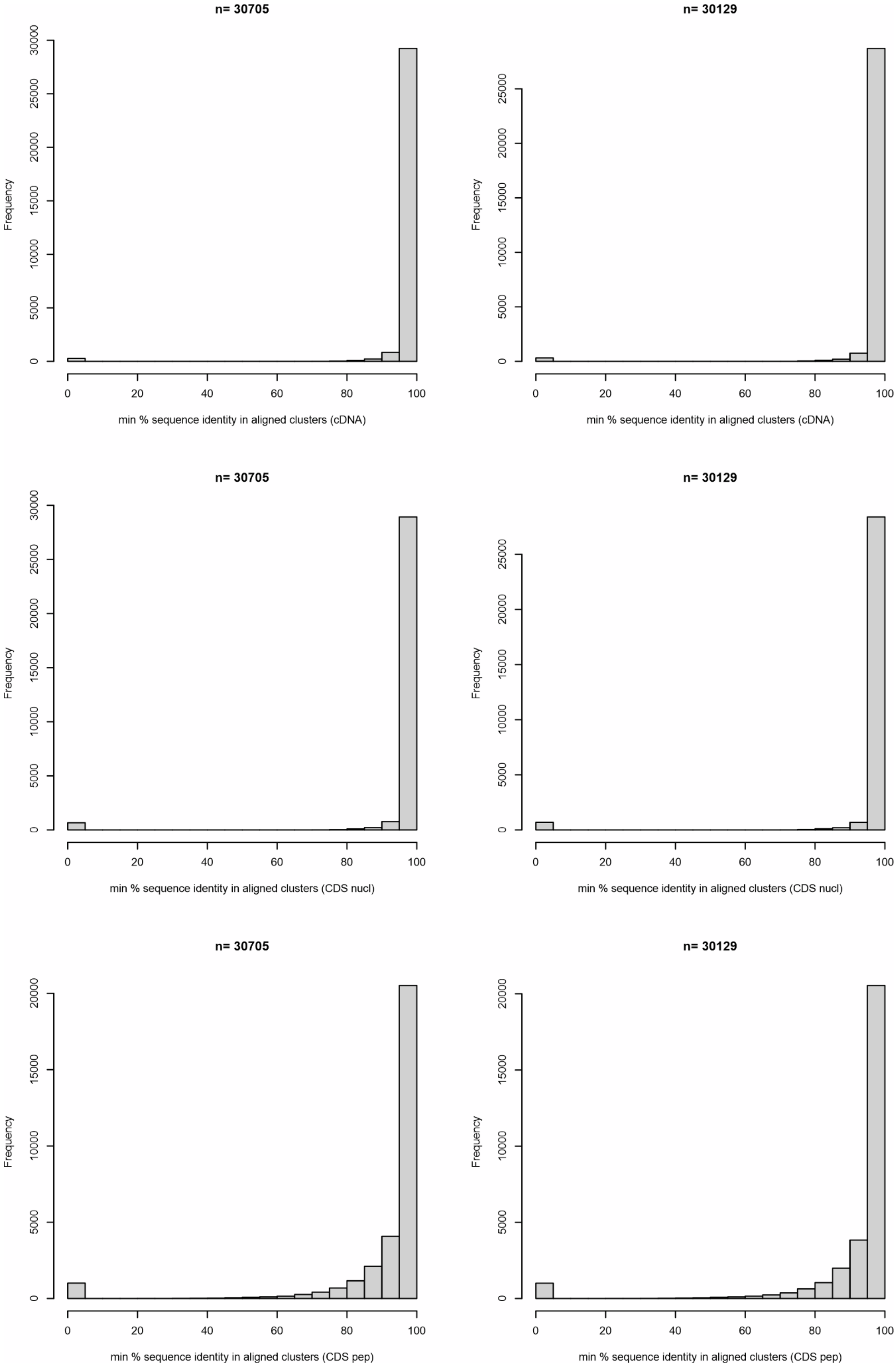
n= 30705
n= 30129
Frequency
min % sequence identity in aligned clusters (cDNA)
min % sequence identity in aligned clusters (CDS nucl)
min % sequence identity in aligned clusters (CDS pep)
0
20
40
60
80
100
5000
10000
15000
20000
25000
30000

**Figure S4**. Sequence identity among sequences in rice3 pangene clusters based on minimap2 (left) and GSAlign (right). Isoform sequences in each cluster were aligned locally with BLAST to the longest one and the lowest identity recorded. Top plots are for cDNA sequences, middle for CDS nucleotide sequences and bottom for CDS protein sequences. As these are local alignments, often only parts of the sequences are actually aligned.

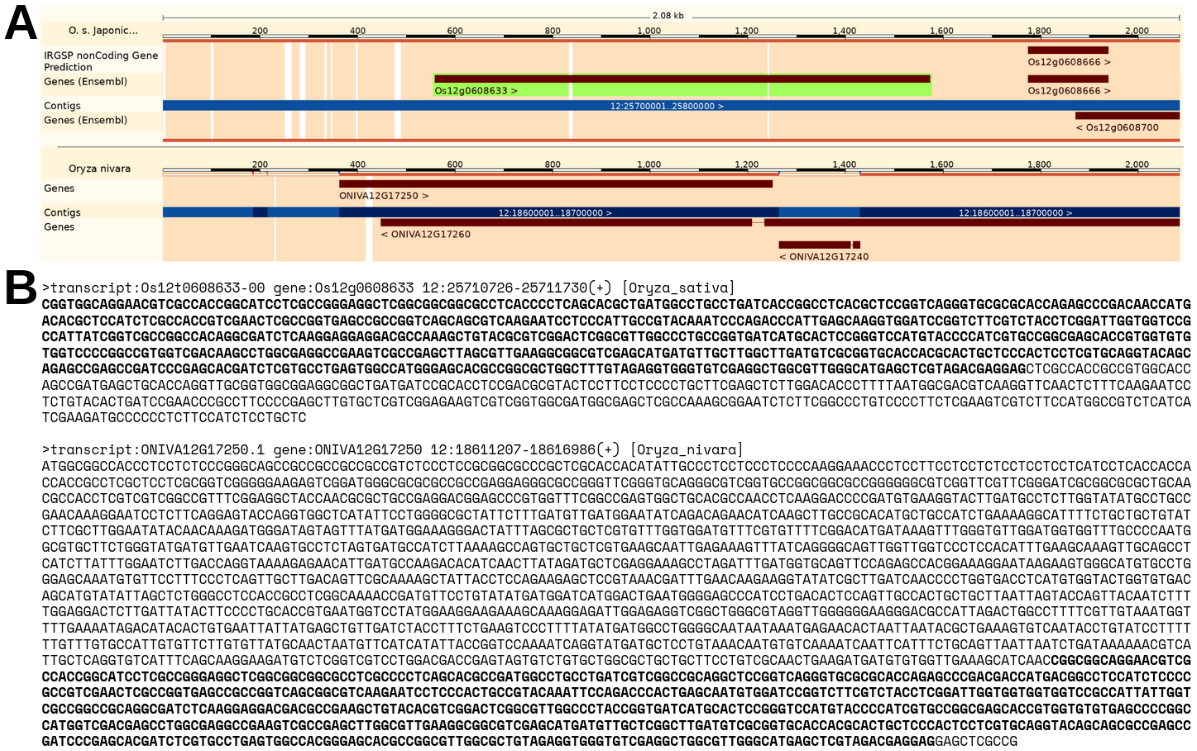


**Figure S5**. Example of pangene cluster where the cDNA sequences have a long local alignment but the encoded CDS sequences cannot be aligned. A) Whole Genome Alignment as displayed in Ensembl Plants browser, where it can be seen that gene models Os12g0608633 (*Oryza sativa* Japonica group, top, green background) and ONIVA12G17250 (*Oryza nivara*, bottom) overlap in the reverse strand. B) cDNA sequences of both gene models, with the shared sequence in bold. In contrast, the CDS sequences of these genes cannot be aligned due to divergent exon annotation.

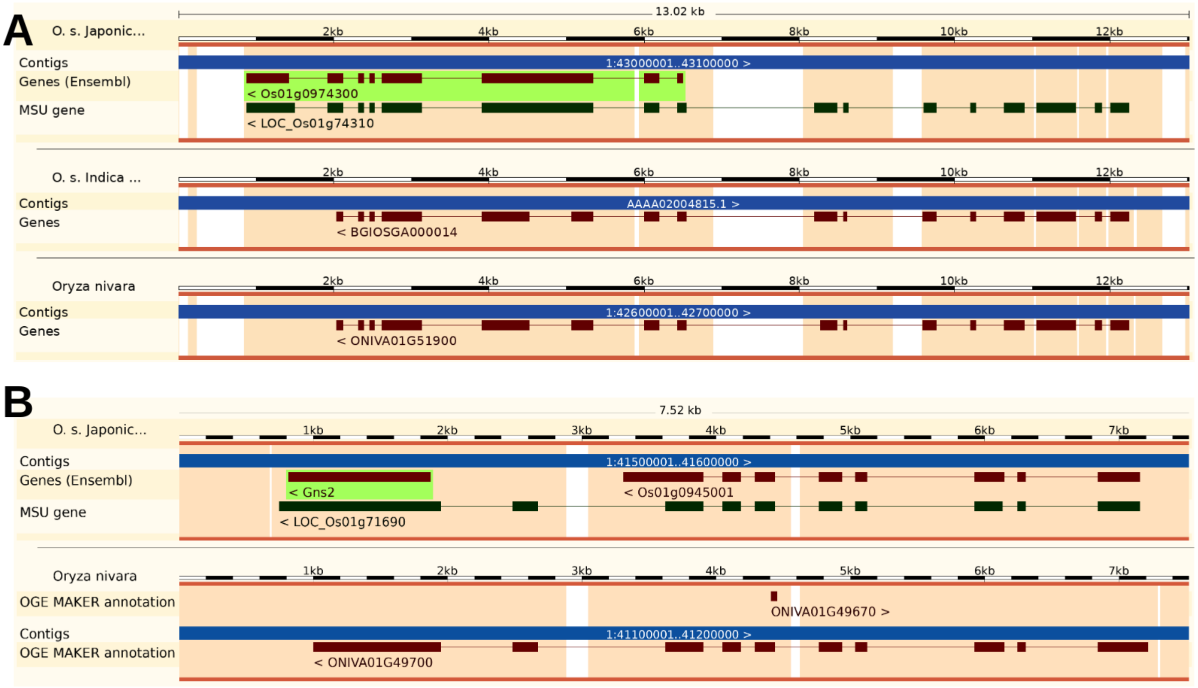


**Figure S6**. Examples of rice pangene clusters not matched by Ensembl Compara orthogroups. In both cases (A, Os01g0974300 and B, Gns2 / Os01g0944900), an *Oryza sativa* Japonica group gene model is split or cut short in the default gene annotation for rice (RAP-DB) (Sakai et al., 2013). As pangene clusters are based on overlapping gene models, split models are correctly grouped together. Moreover, in both cases an alternative annotation source (MSU gene) supports extending or merging the gene models (Kawahara et al., 2013).

A

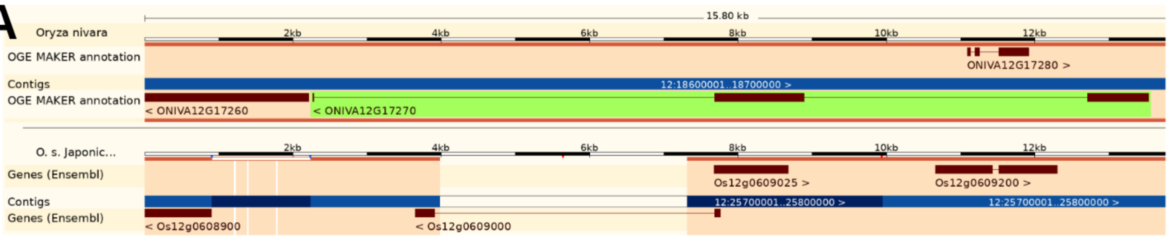


B

>transcript:ONIVA12G17270.1 gene:ONIVA12G17270 12:18626386-18637647(-) [Oryza_nivara]
MQLLFFSFLFLLLARETSAVAADGCSRRCGGLVVPYPFGFSGSCPIMLSCNVDGGSNSTAALILQGNDAT
TTDRSYTVVDGSFNSTASTFTVSVPPSCNRTVSDARRWLSGANYGVSSRTGLFLRGCRNATSSDCSVPVE
TMLRTTRCSGGGGNETASSSLTCIASLSPATPAERGLGGLFAQWEKVEEPRCENLLTSVYGDTREGVFSL
EFAAAEMRWWVNGSCGGGVDDLGRCAANATCIPMQTPSGNWGHRCECLPWMAGDGFAAGEGCYAGKRRRM
RVVEFATAGSVAFLLCLALSVWCLLRRRQWRRNNAKLTVKMARKHLPKDARFFRGKPIEDELELEAAGPR
RFHYGELAAATANFSDDRRLGSGGFGSVYRGFLNGGDVAVKRVAETSRQGWKEFVAEVRIISRLRHRNLV
PLVGWCHDGGDELLLVYELMPNGSLDAHIHSSGNVLPWPARYEVVLGVGAALMYLHHEAEQRVVHRDIKP
SNVMLDASFSARLGDFGLARLIDDGRRSRTTGIAGTMGYIDAECFLLAGRASVESDVYSFGVVLLEVACG
RRPAVVINGGEDAIHLTQWVWDTHGGAAGGGILDAADTRLNGEFDVAEMERVLAVGLWCAHPDRGLRPSI
RQAVSVLRFEAPLPSLPVR**MPVATYGPPVSTASAPTSNDTSAGR**LHP

>transcript:Os12t0609000-00 gene:Os12g0609000 12:25722111-25722886(-) [Oryza_sativa]
**MPVATYGPPVSTASAPTSNDTSAGR**DSATRTVKSEDPLPPRLYARQGQLDSHLFPLAFIEPPFVEHLACM
PIELAIAICLALHLVRRRAPPACACHPLAVLAFLSPWRPSA

**Figure S7**. Example of pangene cluster where the encoded protein sequences do not share protein domains. A) Whole Genome Alignment as displayed in Ensembl Plants browser, where it can be seen that gene models ONIVA12G17270 (*Oryza nivara*, top, green background) and Os12g0609000 (*Oryza sativa* Japonica group, bottom) overlap in the reverse strand. B) Encoded protein sequences by both gene models, with the only shared peptide in bold. This peptide corresponds to the first exon of the *O. sativa* Japonica group and part of the second exon of the *O. nivara*.

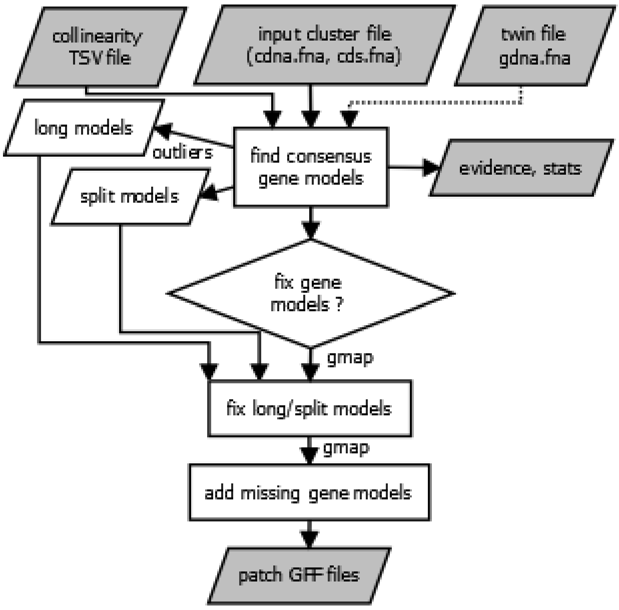


**Figure S8**. Flowchart of script check_evidence.pl , which uses as input a cluster in FASTA format and precomputed collinearity evidence in TSV format. If available, a twin FASTA file with the corresponding collinear genomic segments is also taken, which would be needed for confirming presence/absence of gene models.

```
>Horvu_PLANET_1H01G475600
Paths (1):
  Path 1: query 356..543 (188 bp) => genome 1..188 (188 bp)

            0     .    :    .    :    .    :    .    :    .    :
aa.g        1   F  V  L  F  Q  Q  L  G  R  G  T  V  F  A  P  D
            1 TATTTGTGCTGTTCCAGCAACTAGGCAGGGGTACAGTTTTTGCACCAGAC
              ||||||||||||||||||||||||||||||||||||||||||||||||||
          356 TATTTGTGCTGTTCCAGCAACTAGGCAGGGGTACAGTTTTTGCACCAGAC
aa.c        1   F  V  L  F  Q  Q  L  G  R  G  T  V  F  A  P  D

           50     .    :    .    :    .    :    .    :    .    :
aa.g       17 V  R  Q  N  F  S  C  R  N  F  A  R  Q  Y  H  L  N
           51 GTCCGACAAAACTTCAGCTGCAGGAACTTTGCACGGCAGTACCACCTAAA
              ||||||||||||||||||||||||||||||||||||||||||||||||||
          406 GTCCGACAAAACTTCAGCTGCAGGAACTTTGCACGGCAGTACCACCTAAA
aa.c       17 V  R  Q  N  F  S  C  R  N  F  A  R  Q  Y  H  L  N

          100     .    :    .    :    .    :    .    :    .    :
aa.g       34  V  V  A  A  S  Y  F  N  C  Q  R  E  G  G  S  G  G
          101 CGTTGTGGCTGCCTCATATTTCAACTGTCAAAGGGAAGGTGGATCAGGCG
              ||||||||||||||||||||||||||||||||||||||||||||||||||
          456 CGTTGTGGCTGCCTCATATTTCAACTGTCAAAGGGAAGGTGGATCAGGCG
aa.c       34  V  V  A  A  S  Y  F  N  C  Q  R  E  G  G  S  G  G

          150     .    :    .    :    .    :    .
aa.g       51   R  R  F  R  P  E  S  S  Q  G  E  *
          151 GAAGAAGGTTTAGGCCAGAAAGTTCTCAAGGGGAGTAG
              ||||||||||||||||||||||||||||||||||||||
          506 GAAGAAGGTTTAGGCCAGAAAGTTCTCAAGGGGAGTAG
aa.c       51   R  R  F  R  P  E  S  S  Q  G  E  *
```

**Figure S9**. Partial deletion of locus HvFT3/Ppd-H2 in barley cultivar Igri. A CDS nucleotide sequence encoded by gene Horvu_GOLDEN_1H01G421600 (aa.c) was lifted-over with check_evidence.pl -d MorexV3_highrep_0taxa_5neigh_algMmap_split_ -i gene:HORVU.MOREX.r3.1HG0077240.cds.fna -f -v -n. The alignment against the Igri genome sequence (aa.g) is shown, as computed by GMAP (Wu & Watanabe, 2005). There is a perfect match for nucleotides 356 to 543 of the CDS sequence, which correspond to the last exon (4) of the wild type protein, comprising 61 amino acid residues. Exons 1-3 are not found.

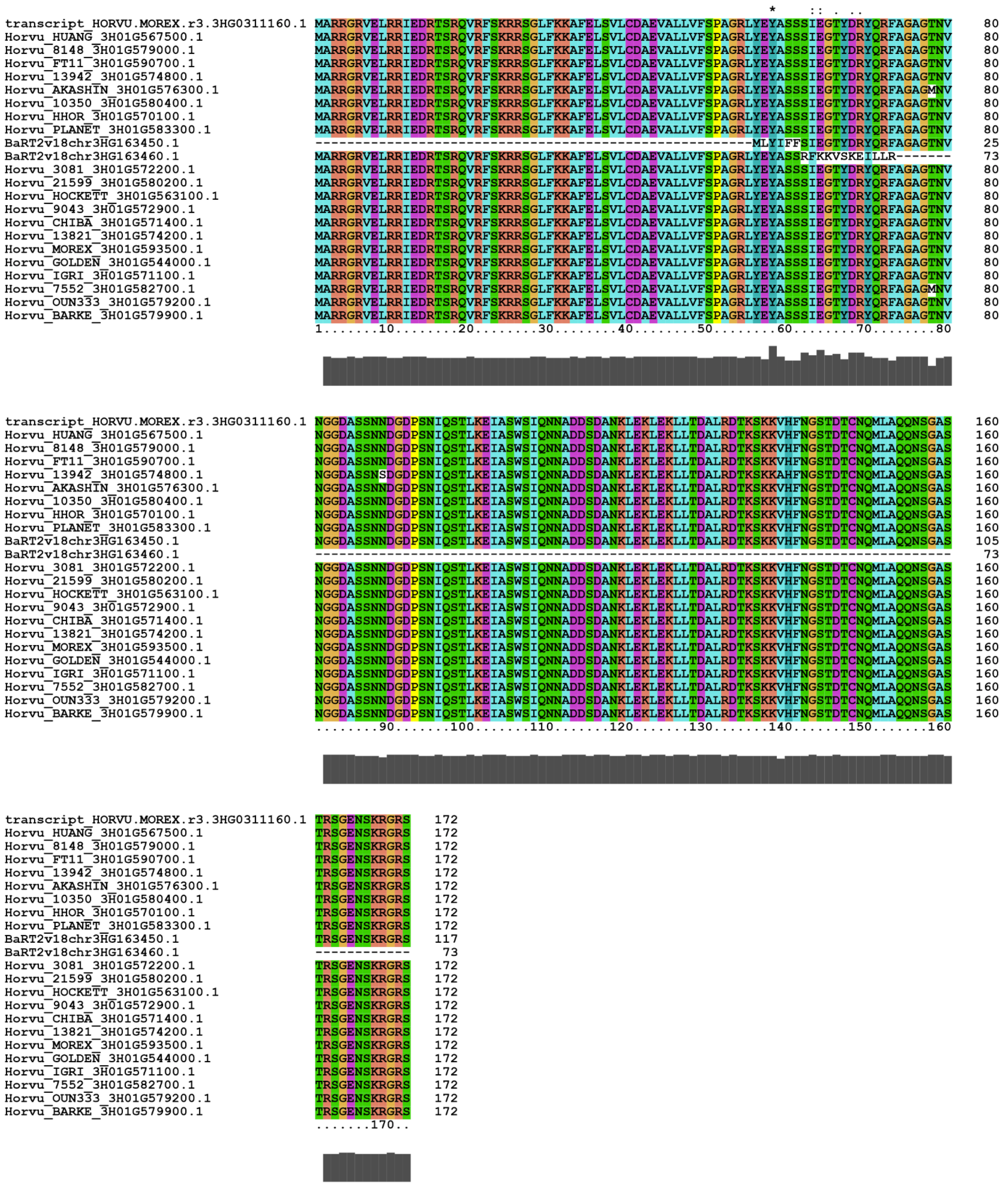

**Figure S10**. Multiple alignment of protein sequences of pangene cluster HORVU.MOREX.r3.3HG0311160, which corresponds to barley locus HvOS2. Isoforms aligned were manually selected and aligned with Clustalx (Larkin et al., 2007).
